# Lysozyme resistance in *C. difficile* is dependent on two peptidoglycan deacetylases

**DOI:** 10.1101/2020.07.17.209676

**Authors:** Gabriela M. Kaus, Lindsey F. Snyder, Ute Müh, Matthew J. Flores, David L. Popham, Craig D. Ellermeier

## Abstract

*Clostridioides (Clostridium) difficile* is a major cause of hospital-acquired infections leading to antibiotic-associated diarrhea. *C. difficile* exhibits a very high level of resistance to lysozyme. Bacteria commonly resist lysozyme through modification of the cell wall. In *C. difficile* σ^V^ is required for lysozyme resistance and σ^V^ is activated in response to lysozyme. Once activated σ^V^, encoded by *csfV*, directs transcription of genes necessary for lysozyme resistance. Here we analyze the contribution of individual genes in the *csfV* regulon to lysozyme resistance. Using CRISPR-Cas9 mediated mutagenesis we constructed in-frame deletions of single genes in the *csfV* operon. We find *pdaV*, which encodes a peptidoglycan deacetylase, is partially responsible for lysozyme resistance. We then performed CRISPR inhibition (CRISPRi) to identify a second peptidoglycan deacetylase, *pgdA*, that is important for lysozyme resistance. Deletion of either *pgdA* or *pdaV* resulted in modest decreases in lysozyme resistance. However, deletion of both *pgdA* and *pdaV* resulted in a 1000-fold decrease in lysozyme resistance. Further, muropeptide analysis revealed loss of either PgdA or PdaV had modest effects on peptidoglycan deacetylation but loss of both PgdA and PdaV resulted in almost complete loss of peptidoglycan deacetylation. This suggests that PgdA and PdaV are redundant peptidoglycan deacetylases. We also use CRISPRi to compare other lysozyme resistance mechanisms and conclude that peptidoglycan deacetylation is the major mechanism of lysozyme resistance in *C. difficile*.

**Importance:** *Clostridioides difficile* is the leading cause of hospital-acquired diarrhea. *C. difficile* is highly resistant to lysozyme. We previously showed that the *csfV* operon is required for lysozyme resistance. Here we use CRISPR-Cas9 mediated mutagenesis and CRISPRi knockdown to show that peptidoglycan deacetylation is necessary for lysozyme resistance and is the major lysozyme resistance mechanism in *C. difficile*. We show that two peptidoglycan deacetylases in *C. difficile* are partially redundant and are required for lysozyme resistance. PgdA provides an intrinsic level of deacetylation and PdaV, encoded as part of the *csfV* operon, provides lysozyme-induced peptidoglycan deacetylation.

## Introduction

*Clostridioides difficile* is a Gram-positive, anaerobic, opportunistic pathogen. *C. difficile* infections are the most common cause of hospital-acquired diarrhea worldwide, with disease severity ranging from mild diarrhea to severe cases of pseudomembranous colitis (1,2). *C. difficile* infections commonly affect patients whose normal gut microbiota has been perturbed by antibiotic treatment. Disease is primarily mediated through two exotoxins, TcdA and TcdB (3, 4). Both toxins are glucosyltransferases that glucosylate Rho family GTPases, leading to cytoskeletal defects in the cell and collapse of tight junctions, which in turn leads to inflammation and cell death of colonic epithelial cells, ultimately resulting in gastrointestinal distress (5, 6).

The bacterial cell wall provides the structure and protection needed for survival in the host environment. In *C. difficile*, the cell envelope is composed of a thick layer of peptidoglycan, a crystalline S-layer, and multiple polysaccharides attached to the cell wall (PS-II and PS-III) (7–10). Presumably to cause an infection *C. difficile* must survive the host immune factors present in the colon and elevated during the inflammatory response. One abundant host defense factor is lysozyme, a component of the innate immune system that cleaves the β-1,4-glycosidic linkage between the GlcNAc and MurNAc residues in the peptidoglycan backbone leading to lysis and cell death (11, 12). *C. difficile* peptidoglycan has unusual features including a high level of GlcNAc *N*-deacetylation and an abundance of 3-3 peptide cross-links (13). Deacetylation of peptidoglycan is a common lysozyme resistance mechanism in a wide variety of bacteria (14).

*C. difficile* senses and responds to lysozyme via the ECF σ factor σ^V^ (15, 16). σ^V^ homologs are also found in multiple other Firmicutes including *Bacillus subtilis* and *Enterococcus faecalis* (17–19). The activity of σ^V^ is inhibited by RsiV, a membrane bound anti-σ factor. In *B. subtilis* and *E. faecalis*, RsiV is degraded in the presence of lysozyme (20, 21). In *C. difficile*, σ^V^ is encoded by *csfV*, which is found in a 7-gene operon (Fig. S1). The σ^V^ operon also encodes *pdaV*, a peptidoglycan deacetylase and *lbpA (cdr20291_1409)*, an *rsiV* ortholog containing a putative lysozyme-binding domain (15).

In order to survive and cause infection, many bacteria modify their cell wall components to resist lysozyme and other cell wall stressors. One mechanism used by multiple organisms including *C. difficile* is alteration of the net charge of the cell envelope through D-alanyl esterification (22–24). A second common resistance mechanism is direct inhibition of lysozyme via lysozyme inhibitor proteins (25–28). The *C. difficile* σ^V^ regulon includes two putative lysozyme inhibitor proteins (RsiV and LbpA) and the *dltABCD* operon which is responsible for D-alanyl esterification (15, 24). Lastly, many bacteria modify their peptidoglycan backbone to prevent lysozyme from accessing the β-1,4-glycosidic linkage (29, 30).

The peptidoglycan backbone can be modified in four ways including acetylation and deacetylation of both the GlcNAc or MurNAc residues (29). *B. subtilis, E. faecalis, Staphylococcus aureus* and *Neisseria gonorrhoeae* resist killing by lysozyme through addition of an acetyl group at the C-6 position of the MurNAc residue, by an O-acetyltransferase frequently encoded by *oatA* (14, 29, 31). *Lactobacillus plantarum* has been shown to have an O-acetyltransferase that modifies the GlcNAc residue (32). Alternatively, other organisms including *B. cereus, B. anthracis, E. faecalis* and *Streptococcus pneumoniae* utilize polysaccharide deacetylases to remove the N-acetyl group on the GlcNAc residue at the C-2 position (14, 29). In *B. subtilis* PdaC has been shown to remove the acetyl group on the MurNAc residue (33). The σ^V^ regulon in *C. difficile* and *E. faecalis* contains a polysaccharide deacetylase (*pdaV* and *pgdA*, respectively) that deacetylates the GlcNAc resides (15, 34). In *B. subtilis* the σ^V^ operon includes *oatA* leading to O-acetylation (19).

Previously, we reported that σ^V^ is required for lysozyme resistance in *C. difficile* and this was partially mediated by deacetylation of peptidoglycan (15). Here we report that the high level of lysozyme resistance and peptidoglycan deacetylation in *C. difficile* is mediated in large part by two peptidoglycan deacetylases, PdaV and PgdA. PgdA provides intrinsic basal deacetylation while PdaV is induced in response to lysozyme, increasing the overall level of deacetylation.

## Results

### σ^V^ is required for expression of the *csfV* operon

σ^V^, encoded by *csfV (sigV)*, in *C. difficile* is necessary for lysozyme resistance (15). Previous studies of *csfV* were performed in a *C. difficile* CD630 derivative, JIR8094, and used a Targetron insertion in *csfV* which is polar on downstream genes (15). Because Targetron mutagenesis does not allow dissecting the role of individual genes, we sought to use CRISPR-Cas9 mutagenesis to construct in frame deletions of genes within the *csfV* operon to determine their contributions to lysozyme resistance. Since σ^V^ is required for its own expression, disruption of *csfV* blocked expression of the entire operon (15, 35, 36). To confirm that σ^V^ is required for expression of the *csfV* operon in *C. difficile* strain R20291, we used CRISPR-Cas9 mediated mutagenesis to construct an in-frame deletion of *csfV* (37). To monitor σ^V^ activity, we used a P_*pdaV*_::*rfp* reporter plasmid to measure activation of the *csfV* operon in response to lysozyme (35). Cultures were grown to mid-log phase, then incubated with lysozyme to induce expression of P_*pdaV*_. In the wildtype strain, we observed a lysozyme-dependent increase in fluorescence indicating increased expression of the P_*pdaV*_::*rfp* reporter (Fig. 1). Consistent with previous studies, lysozyme did not induce P_*pdaV*_::*rfp* expression in the Δ*csfV* mutant (Fig. 1). In fact, basal levels of P_*pdaV*_::*rfp* expression were significantly (~80-fold) lower in the Δ*csfV* mutant, indicating that σ^V^ is required for expression of the *csfV* operon and there is a high basal level expression of the *csfV* operon (15, 35).

**Figure 1.**
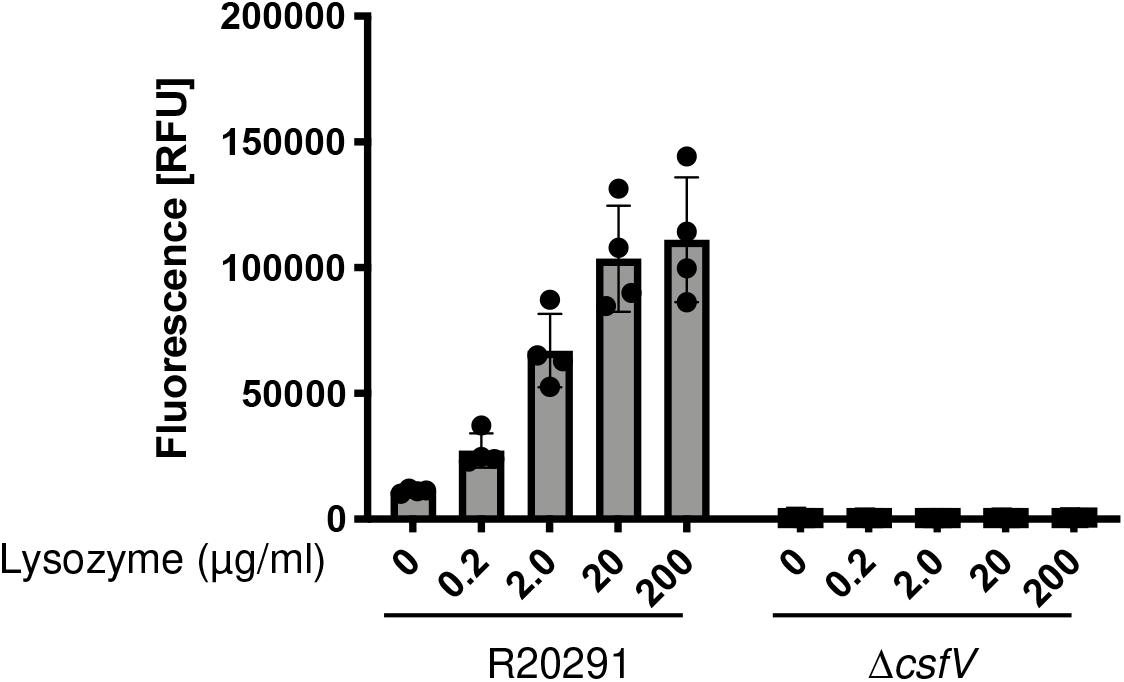
Wildtype (GMK208) or Δ*csfV* (GMK211) strains containing a P_*pdaV*_-*rfp* reporter plasmid were grown to an OD_600_ of 0.3, incubated with lysozyme for 1 hr, then fixed and removed from the anaerobic chamber. Samples were exposed to air overnight to allow for maturation of the chromophore. Fluorescence was measured via a plate reader.

### Activation of σ^V^ increases lysozyme resistance

We developed a liquid culture, 96-well format MIC assay for measuring *C. difficile* sensitivity to lysozyme. Unfortunately, high concentrations of lysozyme (>1 mg/mL) cause turbidity in the medium, preventing a direct read of culture growth. Instead, we evaluated growth by spotting an aliquot on to an agar plate and incubating overnight. Wildtype *C. difficile* is highly resistant to lysozyme with an MIC of 8 mg/ml. We found that the *△csfV* mutant was 4-fold more sensitive to lysozyme than the wildtype (Fig. 2A).

**Figure 2.**
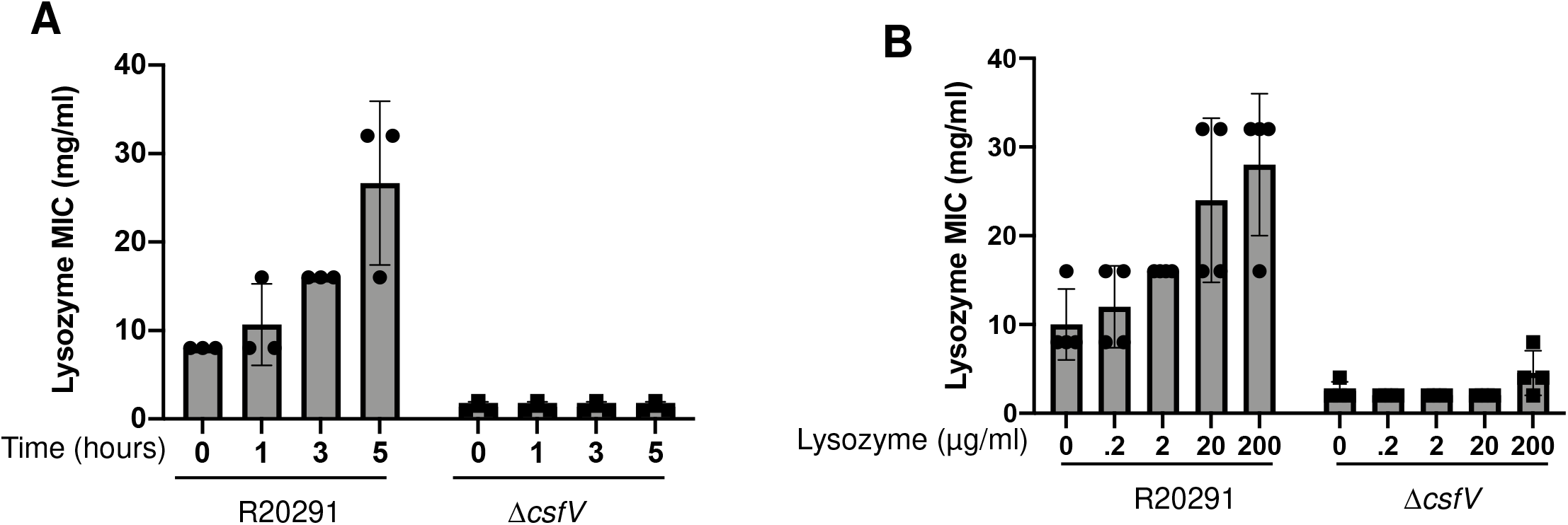
Pre-incubation with a sub-inhibitory concentration of lysozyme increases resistance level. A) Overnight cultures of wildtype or Δ*csfV* (CDE2966) were sub-cultured and grown for ~8hrs; 20 μg/ml lysozyme was added for the duration indicated prior to set up of the lysozyme MIC plates. B) Overnight cultures were sub-cultured and grown to an OD_600_= 0.3; varying sub-inhibitory concentrations of lysozyme were added as indicated and cultures incubated for ~5 hrs prior to set up of MIC.

This MIC assay tests the ability of *C. difficile* to survive exposure to a specific concentration of lysozyme. However, during an infection *C. difficile* likely encounters a gradient of lysozyme concentrations. To reproduce a similar environment, we pre-incubated cultures with a range of sub-inhibitory concentrations of lysozyme prior to exposing them to high levels of lysozyme in the MIC assay. We observed that when incubated with sub-inhibitory lysozyme concentrations, the MIC of wildtype *C. difficile* increases in response to both the concentration and length of exposure (Fig. 2). We found that exposure of wildtype *C. difficile* to 20 μg/ml of lysozyme, for 3 and 5 hours increased lysozyme resistance 2- and 4-fold respectively (Fig. 2A). When the concentration of lysozyme in the pre-incubation step was altered, we saw a dose-dependent increase in the resistance level of the wildtype strain (Fig. 2B). However, we did not observe a change in the MIC of the Δ*csfV* mutant with either increased incubation time or increased lysozyme concentrations, indicating that σ^V^ is required for inducible lysozyme resistance (Fig. 2). We found that pre-incubation for 5 hours with 20 μg/ml lysozyme led to maximum lysozyme resistance in a wildtype with an MIC of 32 mg/ml, which was 16-fold higher than the Δ*csfV* mutant MIC of 2 mg/ml (Fig. 2). A similar observation was made with the Δ*csfV* operon mutant (Fig. S2A).

### PdaV and PgdA are redundant

σ^V^ is required for transcription of genes necessary for lysozyme resistance (15). We sought to dissect the contribution of individual genes to lysozyme resistance. Lysozyme resistance levels were determined using the MIC assay described above. When the samples were grown without pretreatment, we found that when either *csfV* alone or the full *csfV* operon is deleted, the mutant *C. difficile* is 8-fold more sensitive to lysozyme than the wildtype strain (Fig. 3A).

**Figure 3.**
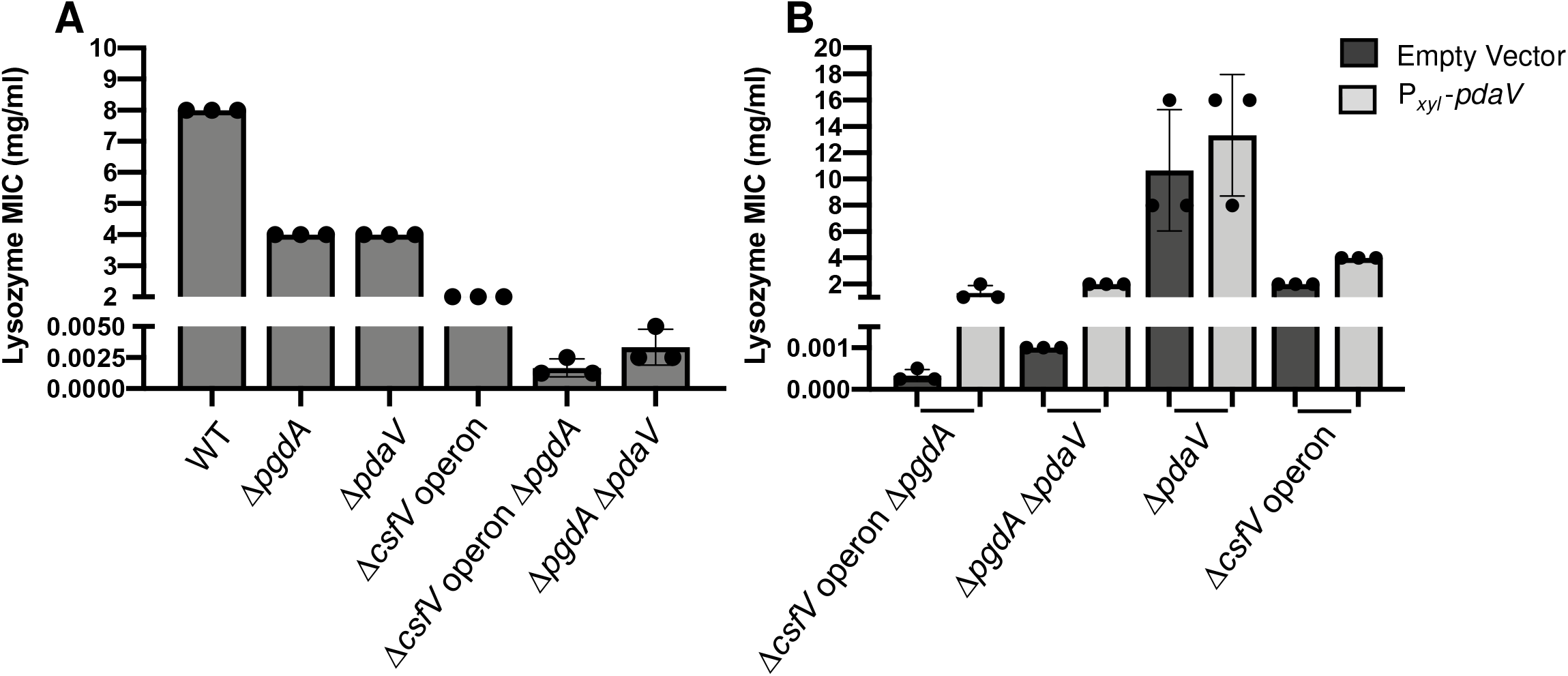
A) Overnight cultures were sub-cultured into TY medium and grown to an OD_600_= 0.3; 20 μg/ml lysozyme was added and incubated for 5 hrs prior to set up of MIC (WT, R20291; Δ*pgdA*, GMK241; Δ*pdaV*, GMK152; Δ*csfV* operon, GMK157; Δ*csfV* operon Δ*pgdA*, GMK243; Δ*pdaV ΔpgdA*, GMK301). B) Strains carrying either P_*xyl*_-*pdaV* (pCE618) or an empty vector (pAP114) were constructed *(ΔcsfV* operon Δ*pgdA* pAP114, GMK312; Δ*csfV* operon Δ*pgdA* pCE618, GMK313; Δ*pdaV ΔpgdA* pAP114, GMK314; Δ*pdaV ΔpgdA* pCE618, GMK315; Δ*pdaV* pAP114, GMK316; Δ*pdaV* pCE618, GMK317; Δ*csfV* operon pAP114, GMK174; Δ*csfV* operon pCE618, GMK177). Overnight cultures were sub-cultured into TY Thi10 medium supplemented with 1% xylose. Cultures were grown to an OD_600_= 1.0 and a lysozyme MIC was set up with 1% xylose.

When *pdaV*, a peptidoglycan deacetylase, is deleted we see a 2-fold reduction in lysozyme sensitivity (Fig. 3A). Additionally, we found that when the Δ*pdaV* mutant was complemented with *pdaV* on a plasmid, lysozyme resistance was restored to levels at or above the wildtype MIC (Fig. 3B). Previous work found that peptidoglycan from a *csfV* mutant in JIR8094 remains highly deacetylated (~75%) (15). Thus, we sought to identify what other factors may be contributing to the high degree of deacetylation. *C. difficile* encodes 7 putative polysaccharide deacetylases that contain predicted transmembrane domains (13, 38). We used CRISPR inhibition (CRISPRi) to knockdown expression of each deacetylase individually and screened for changes in lysozyme sensitivity (39). For each deacetylase gene we constructed two plasmids with different sgRNAs and as a negative control we included a negative control sgRNA that had no target in the *C. difficile* genome. We tested the CRISPRi plasmids in a Δ*csfV* operon strain because we sought to observe differences that are independent of the *csfV* response and it lacked *pdaV* (Fig. S2B). Six of the genes when targeted by CRISPRi had no effect on lysozyme resistance (Fig. S2B). However, we identified one putative deacetylase gene, *cdr20291_1371*, (hereafter referred to as *pgdA* [peptidoglycan deacetylase]) that when knocked down in the Δ*csfV* operon background resulted in a ~100-fold decrease in the level of lysozyme resistance relative to the parent strain (Fig. S2B).

To confirm the results of the CRISPRi knockdown screen, we constructed an in-frame deletion of *pgdA*. When only *pgdA* was deleted we observed a modest 2-fold decrease in lysozyme resistance, similar to the 2-fold decrease observed when *pdaV* is deleted (Fig. 3A). However, when both *pgdA* and *pdaV* are deleted we observed a ~1000-fold decrease in lysozyme resistance (Fig. 3A). This data suggests that *pgdA* and *pdaV* are redundant and that either one is sufficient to confer high level of lysozyme resistance in *C. difficile*. To further support the redundant phenotype of *pgdA* and *pdaV*, we exogenously expressed *pdaV* in the Δ*pgdA ΔpdaV* double mutant. We found that exogenous expression of *pdaV* in a Δ*pgdA ΔpdaV* double mutant restored lysozyme resistance to 2 mg/ml, similar to a Δ*pgdA* mutant (Fig. 3B). We were unable to complement with *pgdA* as we had difficulty cloning in *E. coli* and continuously obtained frameshift mutations within *pgdA*.

Additionally, we used CRISPRi to knockdown expression of *pgdA* in another *C. difficile* strain, JIR8094 an erythromycin sensitive derivative of *C. difficile* CD630 (40). When *pgdA* was knocked down in a wildtype background we observed a ~2-fold reduction in lysozyme resistance (Fig. S3). When *pgdA* was knocked down in a JIR8094 *csfV*-null strain we observed a ~100-fold decrease in lysozyme resistance (Fig. S3). This indicates that loss of both PdaV and PgdA results in a large decrease in lysozyme resistance in multiple *C. difficile* strains.

To determine if expression of *pgdA* was lysozyme-inducible, we constructed a P*pgdA-rfp* fusion. We then tested the effect of increasing lysozyme concentrations on P_*pgdA*_-*rfp* expression. We did not observe an increase in P_*pgdA*_-*rfp* expression in response to increasing lysozyme concentrations suggesting *pgdA* expression is not controlled by lysozyme (Fig. S4). This is consistent with previous microarray experiments that did not show altered expression of *pgdA* by lysozyme (15).

### PdaV and PgdA are the major peptidoglycan deacetylases in *C. difficile*

Next, we sought to identify the acetylation state of peptidoglycan from strains lacking one or both of the deacetylases contributing to lysozyme resistance. We purified peptidoglycan from wildtype, Δ*pgdA*, Δ*pdaV*, and Δ*pgdA* Δ*pdaV* strains. Peptidoglycan was digested with mutanolysin, and muropeptides were separated by charge using reversed-phase HPLC (Fig. 4A). Muropeptide structures were either previously determined or were newly determined using mass spectrometry (15). The peptidoglycan purified from wildtype strain R20291 was made up of muropeptides containing mostly glucosamine (i.e. deacetylated) rather than N-acetylglucosamine residues (Fig. 4B and Table 1). Analysis of the peptidoglycan from the Δ*pdaV* mutant revealed the level of deacetylation remained relatively unchanged compared to the wildtype strain (Fig. 4B and Table 1). When *pgdA* was deleted, we observed decreases in the muropeptides containing glucosamine residues corresponding with an increase in muropeptides containing GlcNAc residues (Fig. 4B and Table 1). In the Δ*pgdA ΔpdaV* mutant we saw a drastic increase in the amount of GlcNAc residues and a concomitant decrease in glucosamine residues, with the overall percentage of muropeptides containing GlcNAc increasing ~15-fold relative to the wildtype strain. For example, we saw in the double mutant the deacetylated tri-gly muropeptides (peak 4) are almost completely absent, concomitant with an increase in acetylated tri-gly muropeptides (peak 3) (Table 1). This suggests that both PdaV and PgdA are major peptidoglycan deacetylases in *C. difficile*.

**Figure 4.**
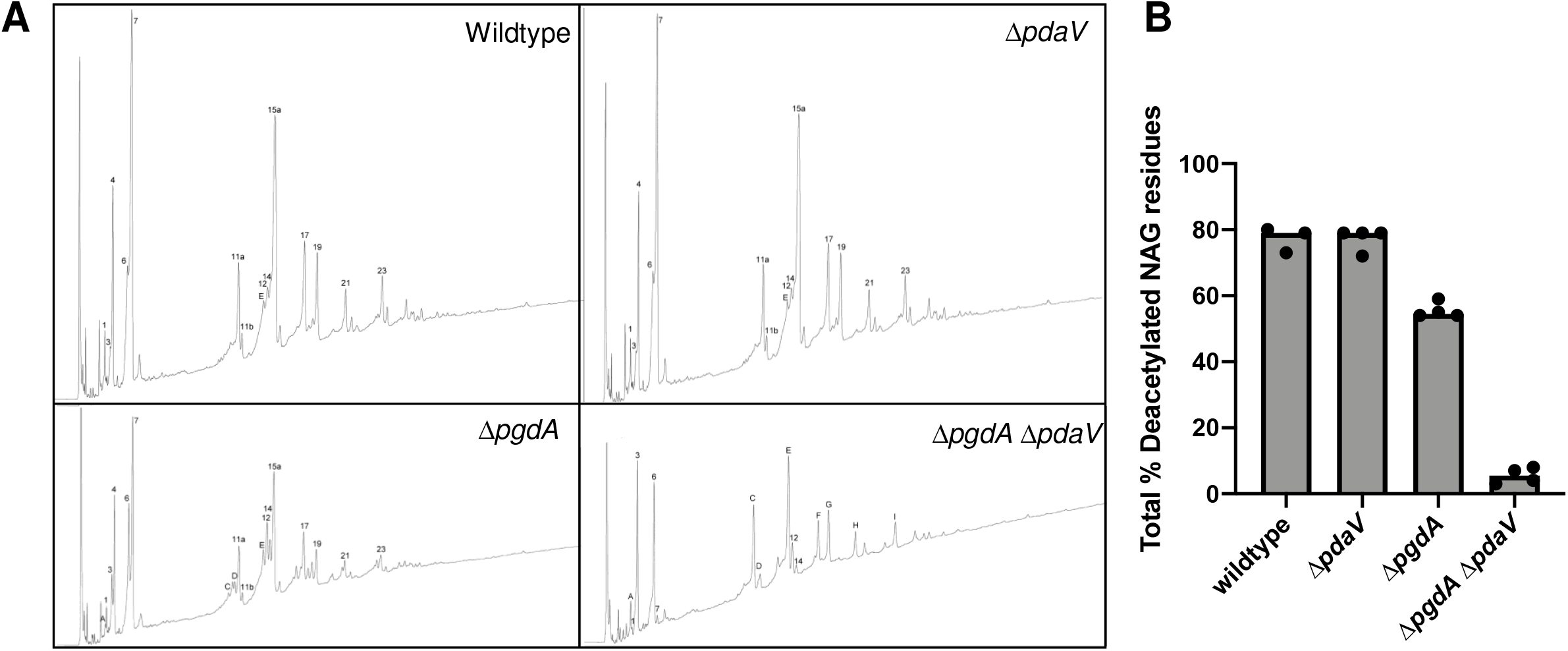
Peptidoglycan was purified from cultures grown to mid-log (OD_600_= 0.6-0.8). Peptidoglycan was digested with mutanolysin. Fragments were separated using reversed-phase HPLC and structures determined using mass spectrometry. B) Total percentages of deacetylated residues are shown for strains indicated (WT, R20291; Δ*pdaV*, GMK152; Δ*pgdA*, GMK241; Δ*pgdA* Δ*pdaV*, GMK301).

**Table 1.**
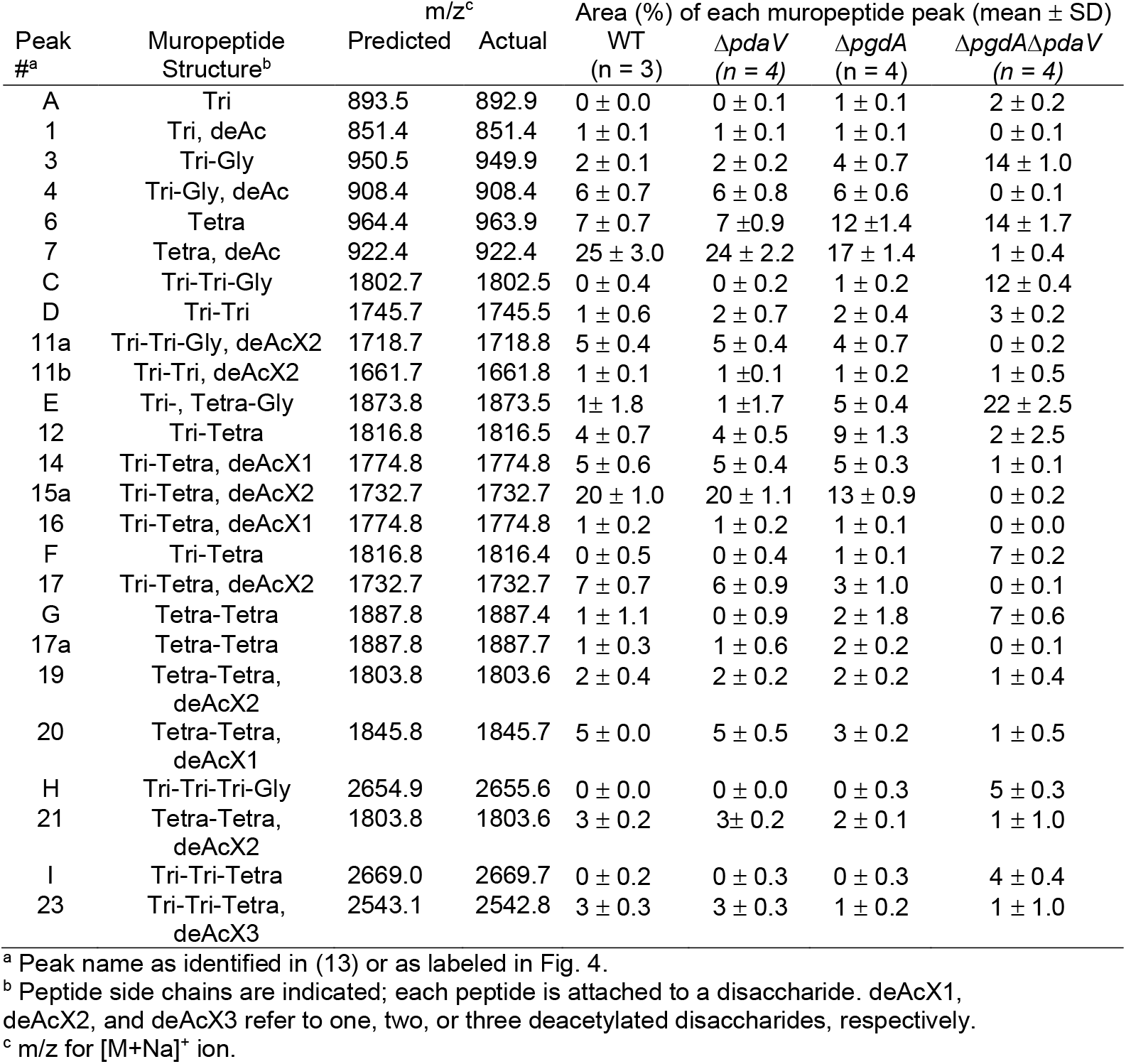
Muropeptide Analysis

| Peak #<sup>a</sup> | Muropeptide Structure<sup>b</sup> | m/z<sup>c</sup> | | Area (%) of each muropeptide peak (mean ± SD) | | | |
| --- | --- | --- | --- | --- | --- | --- | --- |
| | | Predicted | Actual | WT (n = 3) | $\Delta pdaV$ (n = 4) | $\Delta pgdA$ (n = 4) | $\Delta pgdA\Delta pdaV$ (n = 4) |
| A | Tri | 893.5 | 892.9 | 0 ± 0.0 | 0 ± 0.1 | 1 ± 0.1 | 2 ± 0.2 |
| 1 | Tri, deAc | 851.4 | 851.4 | 1 ± 0.1 | 1 ± 0.1 | 1 ± 0.1 | 0 ± 0.1 |
| 3 | Tri-Gly | 950.5 | 949.9 | 2 ± 0.1 | 2 ± 0.2 | 4 ± 0.7 | 14 ± 1.0 |
| 4 | Tri-Gly, deAc | 908.4 | 908.4 | 6 ± 0.7 | 6 ± 0.8 | 6 ± 0.6 | 0 ± 0.1 |
| 6 | Tetra | 964.4 | 963.9 | 7 ± 0.7 | 7 ± 0.9 | 12 ± 1.4 | 14 ± 1.7 |
| 7 | Tetra, deAc | 922.4 | 922.4 | 25 ± 3.0 | 24 ± 2.2 | 17 ± 1.4 | 1 ± 0.4 |
| C | Tri-Tri-Gly | 1802.7 | 1802.5 | 0 ± 0.4 | 0 ± 0.2 | 1 ± 0.2 | 12 ± 0.4 |
| D | Tri-Tri | 1745.7 | 1745.5 | 1 ± 0.6 | 2 ± 0.7 | 2 ± 0.4 | 3 ± 0.2 |
| 11a | Tri-Tri-Gly, deAcX2 | 1718.7 | 1718.8 | 5 ± 0.4 | 5 ± 0.4 | 4 ± 0.7 | 0 ± 0.2 |
| 11b | Tri-Tri, deAcX2 | 1661.7 | 1661.8 | 1 ± 0.1 | 1 ± 0.1 | 1 ± 0.2 | 1 ± 0.5 |
| E | Tri-, Tetra-Gly | 1873.8 | 1873.5 | 1 ± 1.8 | 1 ± 1.7 | 5 ± 0.4 | 22 ± 2.5 |
| 12 | Tri-Tetra | 1816.8 | 1816.5 | 4 ± 0.7 | 4 ± 0.5 | 9 ± 1.3 | 2 ± 2.5 |
| 14 | Tri-Tetra, deAcX1 | 1774.8 | 1774.8 | 5 ± 0.6 | 5 ± 0.4 | 5 ± 0.3 | 1 ± 0.1 |
| 15a | Tri-Tetra, deAcX2 | 1732.7 | 1732.7 | 20 ± 1.0 | 20 ± 1.1 | 13 ± 0.9 | 0 ± 0.2 |
| 16 | Tri-Tetra, deAcX1 | 1774.8 | 1774.8 | 1 ± 0.2 | 1 ± 0.2 | 1 ± 0.1 | 0 ± 0.0 |
| F | Tri-Tetra | 1816.8 | 1816.4 | 0 ± 0.5 | 0 ± 0.4 | 1 ± 0.1 | 7 ± 0.2 |
| 17 | Tri-Tetra, deAcX2 | 1732.7 | 1732.7 | 7 ± 0.7 | 6 ± 0.9 | 3 ± 1.0 | 0 ± 0.1 |
| G | Tetra-Tetra | 1887.8 | 1887.4 | 1 ± 1.1 | 0 ± 0.9 | 2 ± 1.8 | 7 ± 0.6 |
| 17a | Tetra-Tetra | 1887.8 | 1887.7 | 1 ± 0.3 | 1 ± 0.6 | 2 ± 0.2 | 0 ± 0.1 |
| 19 | Tetra-Tetra, deAcX2 | 1803.8 | 1803.6 | 2 ± 0.4 | 2 ± 0.2 | 2 ± 0.2 | 1 ± 0.4 |
| 20 | Tetra-Tetra, deAcX1 | 1845.8 | 1845.7 | 5 ± 0.0 | 5 ± 0.5 | 3 ± 0.2 | 1 ± 0.5 |
| H | Tri-Tri-Tri-Gly | 2654.9 | 2655.6 | 0 ± 0.0 | 0 ± 0.0 | 0 ± 0.3 | 5 ± 0.3 |
| 21 | Tetra-Tetra, deAcX2 | 1803.8 | 1803.6 | 3 ± 0.2 | 3 ± 0.2 | 2 ± 0.1 | 1 ± 1.0 |
| I | Tri-Tri-Tetra | 2669.0 | 2669.7 | 0 ± 0.2 | 0 ± 0.3 | 0 ± 0.3 | 4 ± 0.4 |
| 23 | Tri-Tri-Tetra, deAcX3 | 2543.1 | 2542.8 | 3 ± 0.3 | 3 ± 0.3 | 1 ± 0.2 | 1 ± 1.0 |

<sup>a</sup> Peak name as identified in (13) or as labeled in Fig. 4.
<sup>b</sup> Peptide side chains are indicated; each peptide is attached to a disaccharide. deAcX1, deAcX2, and deAcX3 refer to one, two, or three deacetylated disaccharides, respectively.
<sup>c</sup> m/z for [M+Na]<sup>+</sup> ion.

### RsiV and LbpA act as inhibitors of lysozyme

In *B. subtilis*, RsiV binding to lysozyme leads to σ^V^ activation (21). In addition to RsiV, the σ^V^ operon in *C. difficile* encodes for CDR1409 (CD1560 in JIR8094), an RsiV ortholog hereafter referred to as *lbpA* for lysozyme-binding protein A. LbpA is 58% identical to *C. difficile* RsiV and contains a putative lysozyme binding domain but lacks the σ-binding domain (Fig. S5). We sought to determine if RsiV and LbpA contribute to lysozyme resistance via direct inhibition of lysozyme. To determine if RsiV and LbpA can inhibit lysozyme we performed an *in vitro* lysozyme inhibition assay. We purified 6xHis-RsiV and 6xHis-LbpA from *E. coli*. Peptidoglycan from *M. lysodeikticus* was incubated with lysozyme and varying concentrations of RsiV or LbpA. We observed that when the ratio of RsiV to lysozyme was greater than 1:1, the activity of lysozyme was inhibited, and the peptidoglycan remained intact (Fig. 5A and Fig. S6). When equimolar ratios of lysozyme and RsiV were used most of the lysozyme activity was blocked (Fig. 5A and Fig. S6). Similarly, LbpA inhibited lysozyme activity when used in excess or equimolar ratio to lysozyme (Fig. 5A and Fig. S6). This suggests that both RsiV and LbpA can bind and inhibit lysozyme activity *in vitro*.

**Figure 5.**
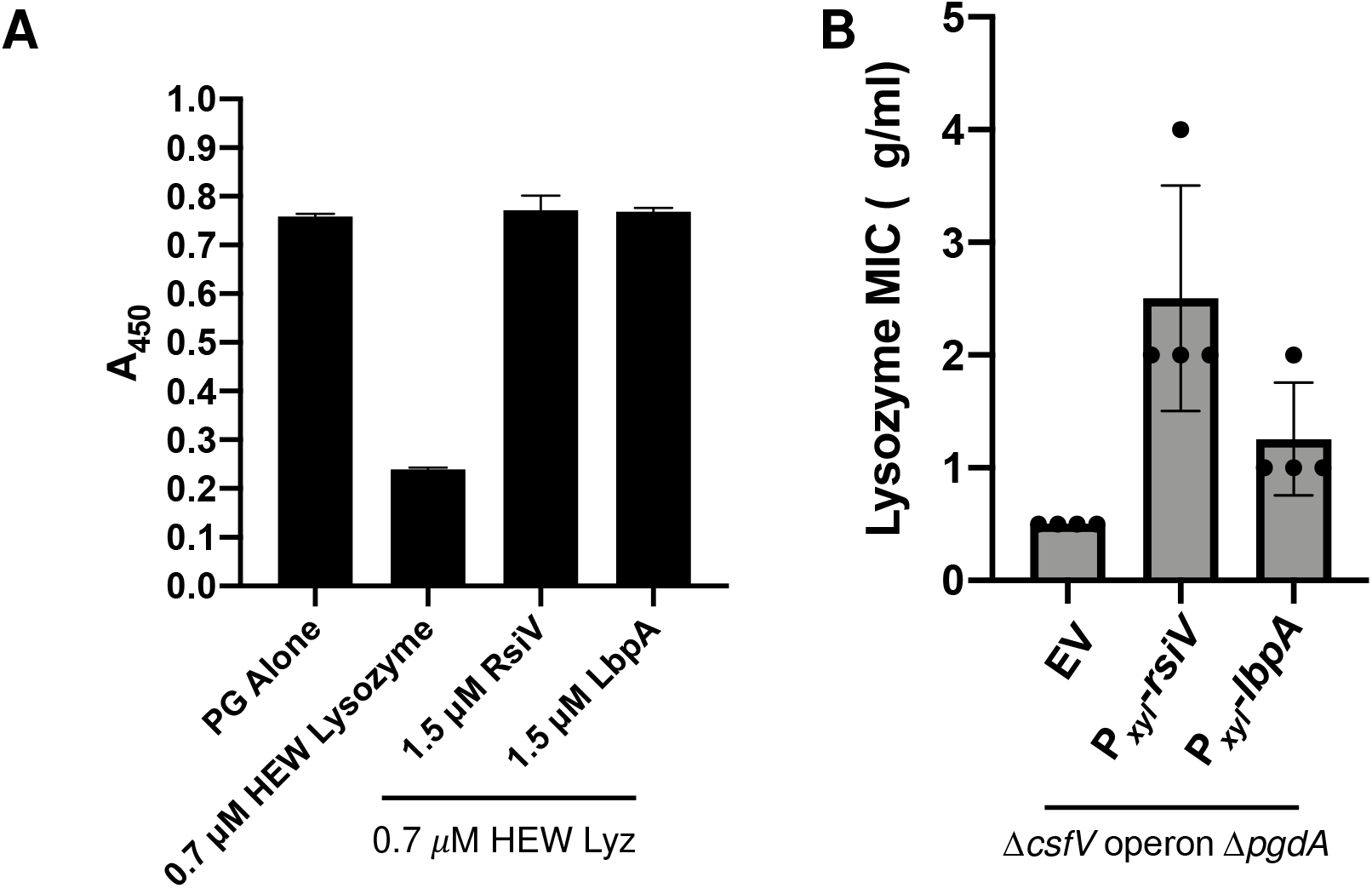
A) Peptidoglycan from *M. lysodeikticus* was combined with 10 μg/ml lysozyme and purified RsiV or LbpA. The A450 was monitored every minute for 30 minutes to determine degradation of lysozyme. Degradation after 30 minutes is shown. B) Overnight cultures were subcultured into TY Thi10 medium supplemented with 1% xylose and grown to OD_600_= 1.0 and a lysozyme MIC was set up with 1% xylose. A one-way ANOVA showed effect of the inhibitor proteins on lysozyme resistance was significant, F (2,9)= 9.8, *p*= 0.0055.

To investigate the effect of LbpA on lysozyme resistance we constructed an in-frame deletion of *lbpA* and tested the effect on the lysozyme MIC. We did not observe any effect on lysozyme resistance (Fig. S7A). We did not attempt to construct a deletion of *rsiV* since it would lead to constitutive activation of σ^V^ and thus would be difficult to dissect the contribution of lysozyme inhibition versus the effect on σ^V^ activity. Instead, to further investigate the role of these putative lysozyme inhibitors we tested the ability of both proteins to alter the resistance level *in vivo* by exogenous expression from the P_*xyl*_ promoter. We found that exogenous expression of *rsiV* or *lbpA* does not alter lysozyme resistance in a Δ*csfV* operon background (Fig. S7B). We hypothesized that inhibition of lysozyme may contribute to lysozyme resistance only when lysozyme concentrations are low. Thus, to detect differences, we used our most lysozyme sensitive strain, Δ*pgdA ΔpdaV*. We observed that exogenous expression of *rsiV* or *lbpA* increases lysozyme resistance 4- and 2-fold respectively, relative to the Δ*pgdA ΔpdaV* parent strain (Fig. 5B). Together these data indicate that both RsiV and LbpA can inhibit lysozyme *in vitro* and confer resistance to low levels of lysozyme *in vivo*.

### Contribution of other cell wall modification to lysozyme resistance

Several other factors have been implicated in lysozyme resistance in *C. difficile* including the S-layer and d-alanylation of teichoic acids (10, 36). We sought to compare the contribution of these other lysozyme resistance mechanisms to the role of peptidoglycan deacetylation. To do this we used CRISPRi to knockdown expression of the *dltABCD* operon, which alters the charge of teichoic acids, and expression of *slpA*, which encodes the major S-layer proteins. We found that when the *dltABCD* operon is knocked down, the MIC decreases ~16-fold from 8 to 0.5 mg/mL (Fig. S8). Similarly, upon knocking down *slpA* we lower the MIC to ~2 mg/ml, a ~4-fold change from wildtype (Fig. S8). As a control for CRISPR-silencing versus gene deletion, we also knocked down the *csfV* operon and observed ~4-8-fold decrease in lysozyme resistance. Thus, knocking down each of these genes results in decreased lysozyme resistance similar to that of the Δ*csfV* mutant. In conclusion, our data suggests that multiple factors contribute to lysozyme resistance. Nevertheless, deacetylation is the single most important mechanism, where deletion of the two key deacetylases achieves an MIC shift of ~1000-fold.

## Discussion

Lysozyme is an important component of the innate immune system and one of the first lines of defense against bacteria. In order to avoid killing by lysozyme, many bacteria modify cell wall properties including peptidoglycan and cell surface charge. We find *C. difficile* encodes both an endogenous lysozyme resistance and an inducible lysozyme resistance mechanism. The inducible lysozyme resistance is primarily mediated by the ECF σ factor σ^V^. In response to lysozyme, σ^V^ upregulates expression of lysozyme resistance mechanisms including the Dlt pathway, a peptidoglycan deacetylase (PdaV), and putative lysozyme inhibitor proteins (RsiV and LbpA) (15, 36). Additionally, a high basal level of peptidoglycan deacetylation and the crystalline S-layer contribute to lysozyme resistance (10). Here we show that together these modifications help confer a high level of lysozyme resistance.

One common mechanism of lysozyme resistance used by bacteria is modification of the acetylation state of peptidoglycan to prevent cleavage at the β-(1-4) linkage by lysozyme. Peptidoglycan modifications include addition or removal of acetyl groups from either the MurNAc or GlucNAc residues on the peptidoglycan backbone (29, 41). In many pathogens, including *S. aureus* and *L. monocytogenes* O-acetylation is important for resistance to lysozyme and virulence (42, 43). Multiple organisms, including *C. difficile, B. subtilis* and *E. faecalis* modify peptidoglycan in a σ^V^ dependent manner in response to lysozyme (15, 18, 19, 31, 44). In *B. subtilis*, the σ^V^ operon includes an O-acetylase, *oatA* which is responsible for acetylating the MurNAc residues in peptidoglycan (19, 31). Additionally, the *dlt* operon in *B. subtilis*, responsible for D-alanylation of teichoic acids, is regulated by σ^V^ (31, 45). Mutations in either *oatA* or the *dlt* operon results in ~2-fold decrease in lysozyme resistance (19, 31). Similarly, the σ^V^ regulon in *E. faecalis* includes *pgdA*, a peptidoglycan deacetylase that contributes to lysozyme resistance (44, 46). *E. faecalis* also depends on OatA and the Dlt pathway for lysozyme resistance, however these are σ^V^-independent (18, 34, 44).

Previous work demonstrated that *C. difficile* peptidoglycan is highly deacetylated (13). We had shown that deacetylation can be increased in response to lysozyme via increased activity of σ^V^ which was presumably mediated by PdaV (15). Here, we have identified an additional peptidoglycan deacetylase, PgdA, that plays a critical role in lysozyme resistance. When both *pgdA* and *pdaV* are deleted, only a small percentage of the peptidoglycan is deacetylated and the level of lysozyme resistance is drastically decreased. This suggests that the high level of deacetylation observed in *C. difficile* is required for lysozyme resistance. Our data indicated either PgdA or PdaV are sufficient for high levels of lysozyme resistance, with loss of either *pgdA* or *pdaV* resulting in modest 2-fold decreases in lysozyme resistance. When *pgdA* alone was deleted, we detected a decrease in total percentage of deacetylated glucosamine residues. Interestingly, we did not observe a change in total percentage of deacetylated residues in the absence of *pdaV*. This is likely because the samples were grown in the absence of lysozyme and therefore the σ^V^ response was not activated. However, it is also clear from this analysis that even in the absence of lysozyme there is a significant contribution of PdaV to deacetylation likely due to high basal level expression of *pdaV*. In fact, we observed high basal level expression of the *pdaV* reporter in a wildtype strain and deletion of *csfV* resulted in a marked decrease in *PpdaV-rfp* reporter signal. Taken together our data suggest that either PdaV or PgdA are sufficient for peptidoglycan deacetylation and lysozyme resistance in *C. difficile*.

We have shown that exposing cultures to sub-inhibitory levels of lysozyme prior to incubation with high levels of lysozyme allows the cells to increase the lysozyme MIC in a σ^V^-dependent manner. This indicates that the modifications made by the σ^V^ regulon during exposure to sub-inhibitory lysozyme permit the bacteria to increase their ability to survive when exposed to high levels of lysozyme. Lysozyme activates the σ^V^-mediated modifications including increased deacetylation and D-alanylation of teichoic acids (36).

One of the unique features of the *C. difficile csfV* operon is the presence of two proteins, RsiV and LbpA, which can bind lysozyme. RsiV functions as an anti-σ factor and can inhibit σ^V^ activity (19). In contrast, LbpA lacks the σ^V^-binding domain. We find that both RsiV and LbpA inhibit lysozyme activity *in vitro*. Our data also show that exogenous production of either RsiV or LbpA in a lysozyme sensitized strain *(ΔpgdA ΔpdaV)* increases lysozyme resistance. The ability of RsiV and LbpA to inhibit lysozyme activity and increase the MIC in a sensitive strain indicate a possible role for lysozyme inhibitors in *C. difficile*. It is unclear how rapidly the peptidoglycan and lipoteichoic acids can be modified to increase lysozyme resistance. As a *C. difficile* infection is established, the toxins elicit an inflammatory response recruiting neutrophils to the site of infection increasing the concentration of lysozyme. We hypothesize that the lysozyme inhibitors may be important early in infection to sequester lysozyme allowing the cell additional time for surface modifications such as increased deacetylation.

## Materials and Methods

### Bacterial strains, media and growth conditions

Bacterial strains are listed in Table 2. *C. difficile* strains used in this study are derivatives of R20291. *C. difficile* was grown in or on tryptone-yeast (TY) medium supplemented as needed with thiamphenicol at 10 μg/ml (Thi10), kanamycin at 50 μg/ml, or cefoxitin at 50 μg/ml. TY consisted of 3% tryptone, 2% yeast extract and 2% agar (for solid medium). *C. difficile* strains were maintained at 37°C in an anaerobic chamber (Coy Laboratory products) in an atmosphere of 10% H2, 5% CO_2_, and 85% N_2_.

**Table 2.**
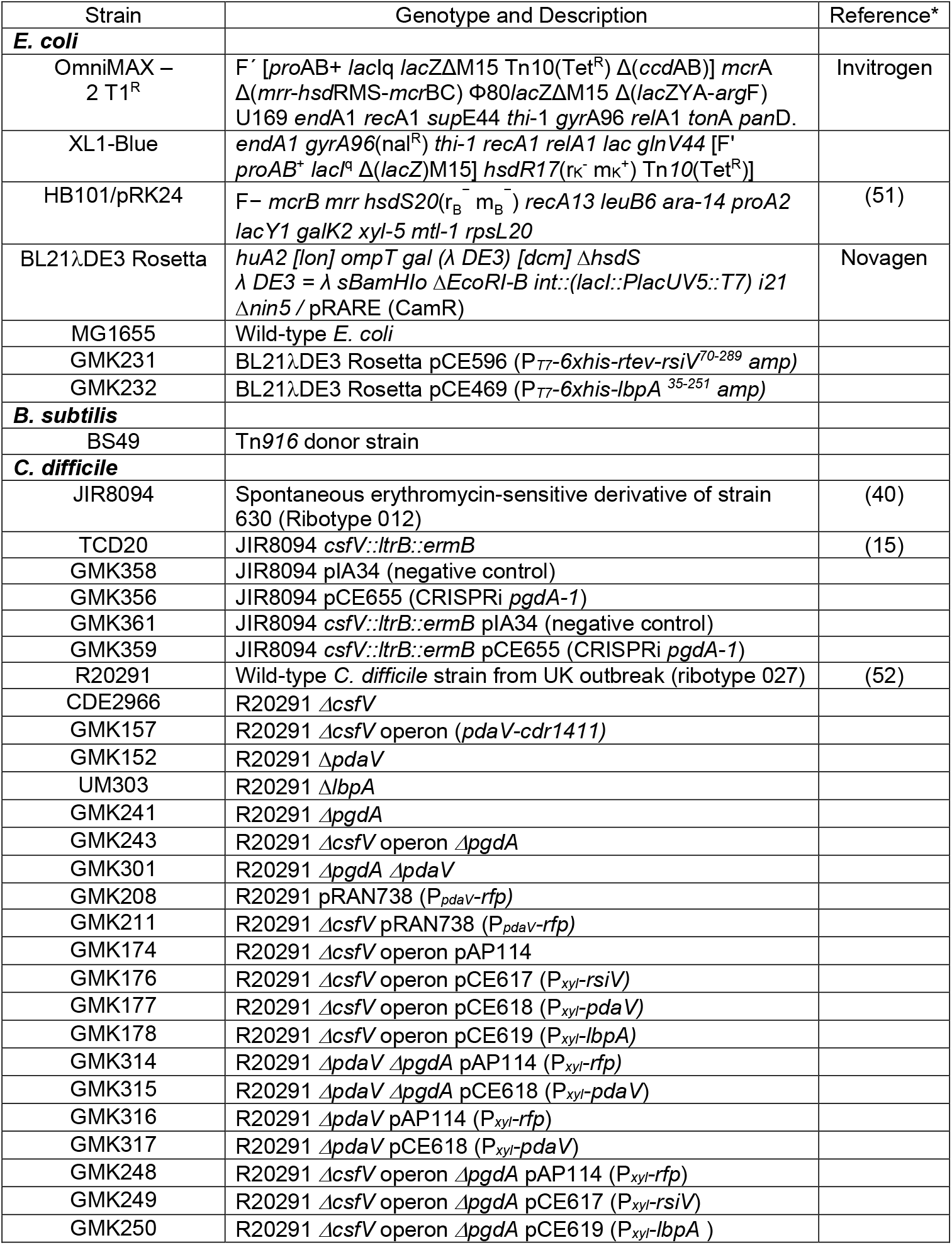
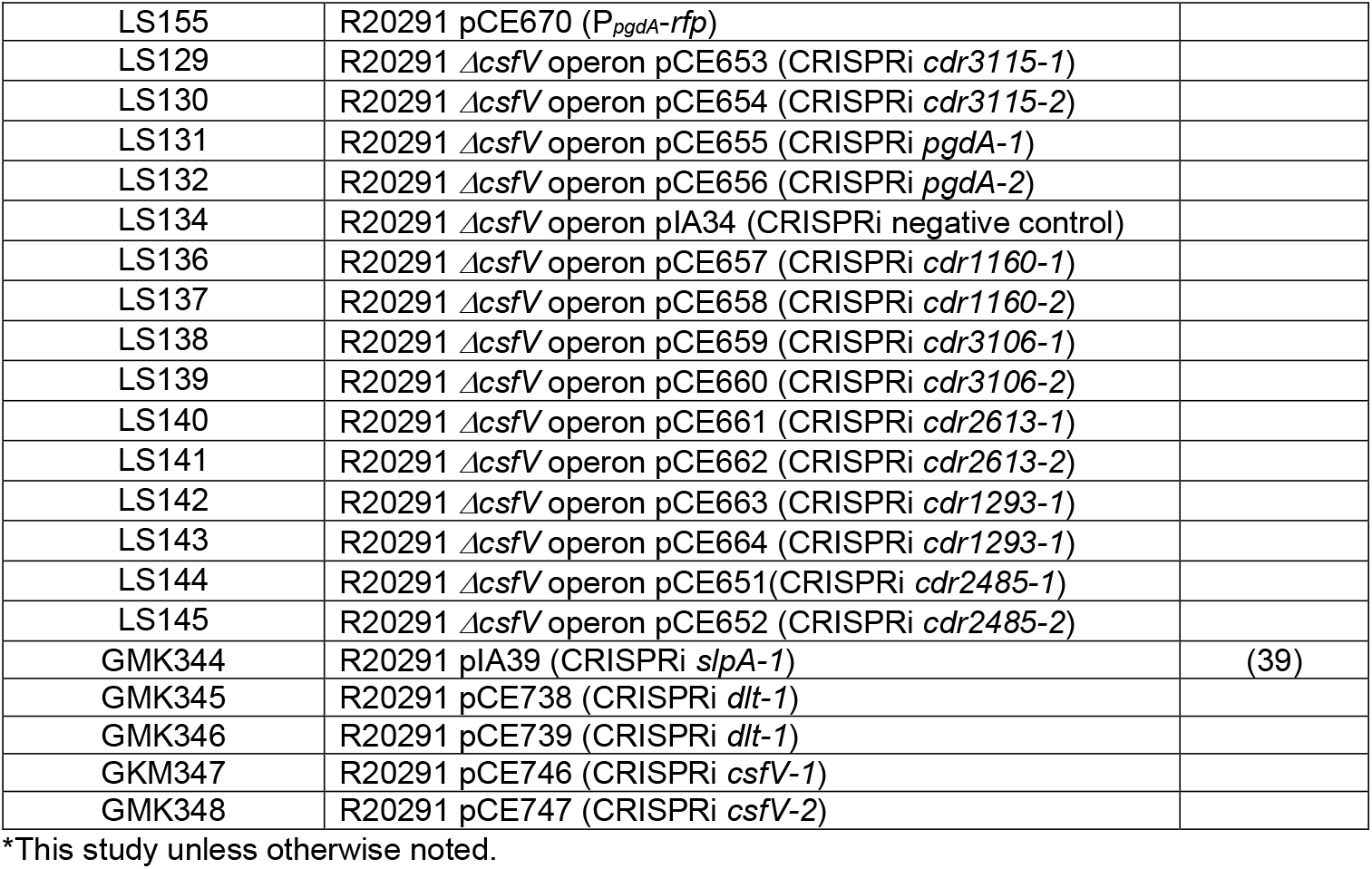
Strains

*E. coli* strains were grown in LB medium at 37°C with chloramphenicol at 10 μg/ml and ampicillin at 100 μg/ml as needed. LB contained 1% tryptone, 0.5% yeast extract, 0.5% NaCl and 1.5% agar (for solid medium).

### Plasmid and bacterial strain construction

All plasmids are listed in Table 3 and Table S1. Plasmids were constructed using Gibson Assembly (New England Biolabs, Ipswich, MA). Regions of plasmids constructed using PCR were verified by DNA sequencing. Oligonucleotide primers used in this work were synthesized by Integrated DNA Technologies (Coralville, IA) and are listed in Table S2. All plasmids were propagated using OmniMax-2 T1R as a cloning host. CRISPR-Cas9 deletion plasmids were passaged through *E. coli* strain MG1655, before transformation into *B. subtilis* strain BS49. The CRISPR-Cas9 deletion plasmids which harbor the *oriT*_(Tn916)_ origin of transfer, were then introduced into *C. difficile* strains by conjugation (37). All other plasmids (RP4 *oriT traJ* origin of transfer) were transformed into *E. coli* strain HB101/pRK24, then introduced into *C. difficile* by conjugation (47).

**Table 3.**
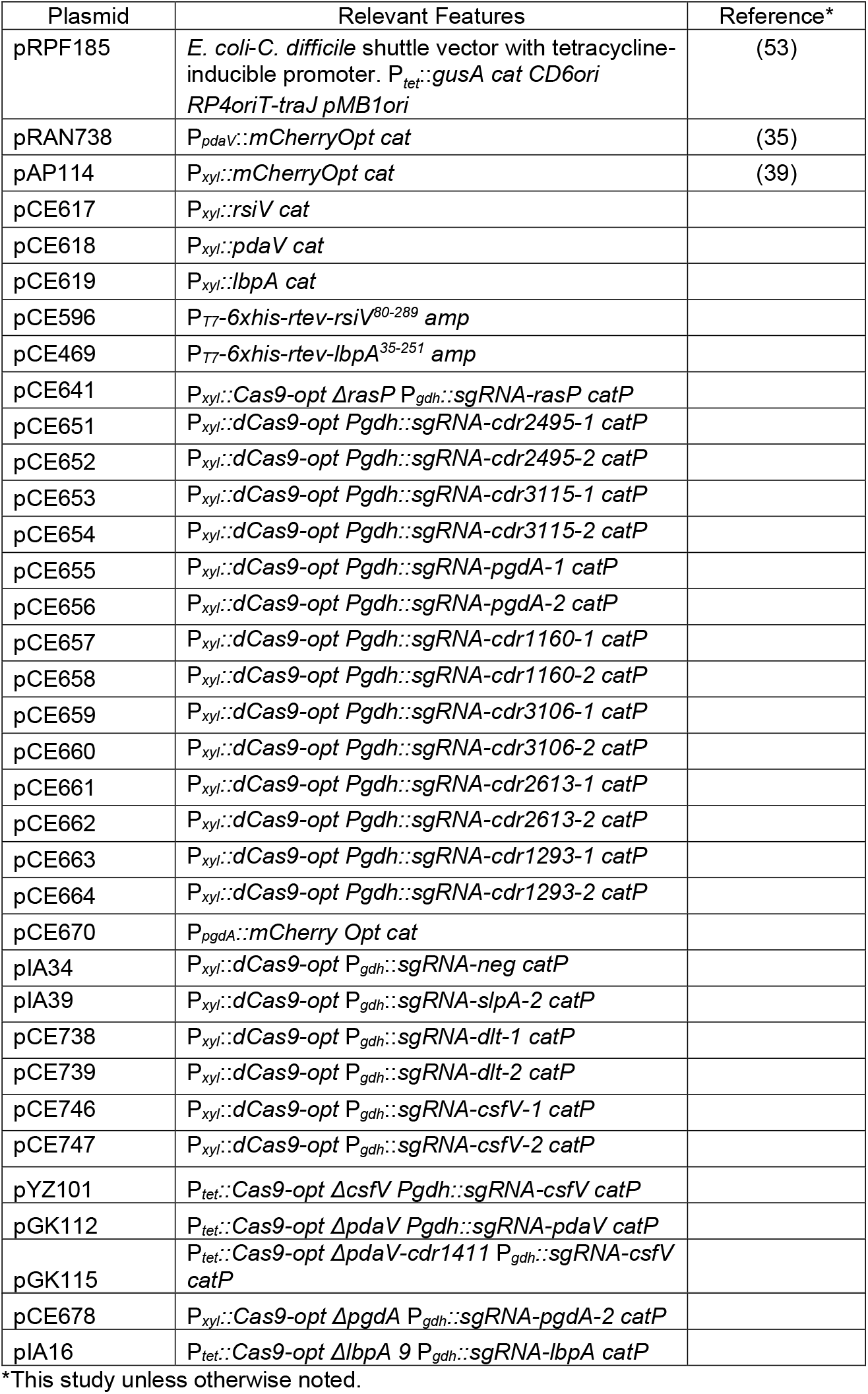
Plasmids

CRISPR-Cas9 plasmids were built on the backbone of pJK02 (37). Initial constructs expressed *cas9* under P_*tet*_ control. The final construct, pCE641, placed *cas9* under P_*xyl*_ control. The P_*tet*_ regulatory element was removed from pJK02 by digestion with PacI and XhoI and replaced with a *xylR-P_xyl_* fragment amplified by PCR of R20291 chromosomal DNA. Donor regions for homology were made by separately amplifying regions ~500 bp upstream and ~500 bp downstream of the gene of interest. The resulting regions were cloned into the NotI and XhoI restriction sites in pCE641 by Gibson Assembly. The algorithm provided by Benchling was used to design sgRNAs targeting each deleted gene (48). Guide parameters were set to default conditions to identify a 20-nucleotide guide with the PAM set to NGG. sgRNA fragments were amplified by PCR from pCE641, using an upstream primer that introduces the altered guide and inserted at the KpnI and MluI sites of the pCE641-derivative with the appropriate homology region.

For CRISPRi constructs, the algorithm provided by Benchling was used to design sgRNAs as described above. Final candidates were selected to be high scoring and bind to the non-coding strand in the first third of the gene sequence. The sequences for sgRNAs are summarized in Table S3. sgRNA fragments were amplified by PCR from pIA33, using an upstream primer that introduces the altered guide and inserted at the MscI and NotI sites of the pIA33 (39).

### Peptidoglycan purification

Peptidoglycan was purified from 100 ml cultures grown to an OD_600_ of 0.6 to 0.8 in TY broth. Peptidoglycan was purified as previously described (Ho 2014). Briefly, cells were pelleted by centrifugation and supernatant was discarded. The cells were boiled in 4% sodium dodecyl sulfate (SDS) for 1 hr. Samples were pelleted and supernatant was discarded. Samples were washed in sterile water 6 times to remove SDS. Samples were digested with DNaseI (NEB) and RNase (Sigma-Aldrich) for 2 hrs at 37°C to remove nucleotides and then digested with Trypsin (Sigma-Aldrich) for 16 hrs at 37°C. Teichoic acids were removed by resuspending the samples in 1 ml 49% hydrofluoric acid (VWR) rocking for 48 hrs 4°C. Samples were subsequently washed extensively in sterile water. The peptidoglycan was digested for 16 hrs with 125 units Mutanolysin (Sigma-Aldrich) as previously described (49). Muropeptides were separated and purified using HPLC and analyzed by MALDI-TOF MS as previously described (15).

### Protein expression and purification

Cultures of *E. coli* BL21λDE3 Rosetta containing an expression plasmid for either RsiV or LbpA were diluted 1:100 in 100 ml LB supplemented with 10 μg/ml chloramphenicol and 100 μg/ml ampicillin. Samples were grown at 30°C to an OD_600_ of 0.6 then induced with 1mM IPTG and grown for an additional 4 hrs at 30°C. Cells were chilled on ice and pelleted by centrifugation at 2500 x *g*. Cell pellets were stored at −80°C until time for purification. Cells were thawed on ice in 2 ml lysis buffer (50 mM Tris HCl 250 mM NaCl, 10 mM imidazole, 3 mM Triton X-100 pH 8.0). Cells were lysed by sonication and lysate was centrifuged at 17,000 x *g* to pellet cell debris. Clarified lysate was added to 500 μl nickel resin slurry (Thermo) and incubated rocking at 4°C for 30 minutes to bind 6xHis-tagged protein. The resin was washed five times with 2 ml wash buffer (50 mM Tris-HCl, 250 mM NaCl, 20 mM imidazole, 0.3 mM Triton X-100 pH 8.0). To elute protein, resin was incubated with 500 μl elution buffer (50 mM Tris-HCl, 250 mM NaCl, 250 mM imidazole, 0.03 mM Triton X-100 pH 8.0) rocking at 4°C for 15 minutes. Samples were centrifuged, and supernatant collected.

### Lysozyme activity assay

Lysozyme activity was measured by following the degradation of a commercially available PG preparation (lyophilized *Micrococcus lysodeikticus* cells from Sigma). Increasing concentrations of purified RsiV or LbpA were mixed with lysozyme (final concentration in assay, 10 μg/ml) in 50 μl in a 96 well plate. The reaction was started by the addition of 50 μl PG substrate (*M. lysodeikticus* PG suspension in 50 mM Na phosphate, pH 7, 100 mM NaCl at an optical density of 1.8). PG lysis was monitored at 450 nm, every minute for 30 min (M200 Pro plate reader, Tecan).

### Lysozyme MIC determination

Overnight cultures were sub-cultured and grown to late log phase (OD_600_ of 1.0), then diluted in TY to 10^6^ CFU/ml. For samples that were pre-incubated with lysozyme, lysozyme was added at the time and concentration indicated. A series of lysozyme concentrations was prepared in a 96-well plate in 50 μl TY broth. Wells were inoculated with 50 μl of the dilute late log culture (i.e. 0.5 x 10^5^ CFU/well) and grown at 37°C for 16 hrs. Each well was then sampled by removing 10 μl and diluting 1:10 in TY broth; 5 μl of this dilution was spotted onto TY agar and incubated at 37°C for 24 hrs. The MIC was defined as the lowest concentration of lysozyme at which 5 or fewer colonies were found per spot.

### Fixation protocol

Cells were fixed as previously described (35, 50). Briefly, a 500 μl aliquot of cells in growth medium was added to 100 μl 16% paraformaldehyde (Alfa Aesar) and 20 μl of 1M NaPO4 buffer (pH 7.4). The sample was mixed, removed from the chamber, and incubated in the dark at room temperature for 60 minutes. The samples were washed 3 times with phosphate-buffered saline (PBS), resuspended in 100 μl PBS, and left in the dark for a minimum of 3 hrs to allow for maturation of the chromophore.

### Fluorescence measurements with a plate reader

Fluorescence from bulk samples was measured using an Infinite M200 Pro plate reader (Tecan, Research Triangle Park, NC) as previously described (35, 39). Briefly, fixed cells in PBS were added to a 96-well microtiter plate (black, flat optical bottom). Fluorescence was recorded as follows: excitation at 554 nm, emission at 610 nm, and gain setting 140. The cell density (OD_600_) was also recorded and used to normalize the fluorescence reading.

## Acknowledgements

This work was supported by the National Institutes of Allergy and Infectious Disease NIH R01AI087834 to CDE and a graduate fellowship T32AI007511 to GMK. We thank Joe Sorg for providing the CRISPR mutagenesis plasmid. We would also like to thank members of the Ellermeier and Weiss labs for helpful comments.

---

**Figure S1.** Organization of the *csfV* operon in *C. difficile* strain R20291.

**Figure S2.** Contribution of putative polysaccharide deacetylases to the lysozyme MIC. A) MICs of in-frame deletions of individual genes (WT, R20291; Δ*csfV*, CDE2966; Δ*csfV* operon, GMK157). Overnight cultures were sub-cultured and grown to an OD_600_ of 0.3, incubated with 20 μg/ml lysozyme for 5 hrs, then MICs were determined. B) Putative polysaccharide deacetylases were screened using CRISPRi knockdown in a Δ*csfV* operon strain *(cdr1160*, LS136, LS137; *cdr1293*, LS142, LS143; *pgdA*, LS131, LS132; *cdr2485*, LS144, LS145; *cdr2613*, LS140, LS141; *cdr3106*, LS138, LS139; *cdr3115*, LS129, LS130; CRISPRi negative control, LS134). Overnight cultures grown with 1% xylose were sub-cultured in TY supplemented with 1% xylose and grown to an OD_600_ of 1.0, then MICs were determined.

**Figure S3.** CRISPRi knockdown of *pgdA* was tested in JIR8094 strain backgrounds (JIR8094 CRISPRi negative control, GMK358; JIR8094 CRISPRi *pgdA*, GMK256; JIR8094 *csfV::ltrB::ermB* CRISPRi negative control, GMK361; JIR8094 *csfV::ltrB::ermB* CRISPRi *pgdA*, GMK359). Overnight cultures grown with 1% xylose were sub-cultured in TY supplemented with 1% xylose and grown to an OD_600_ of 1.0, then MICs were determined.

**Figure S4.** *pgdA* is not activated by lysozyme. Strains carrying either *PpdaV-rfp* (GMK208) or *PpgdA-rfp* (LS155) were sub-cultured and grown to an OD_600_ of 0.3 then incubated with varying concentrations of lysozyme for 1 hr. Cultures were fixed, removed from the chamber and exposed to air overnight to allow for maturation of the chromophore. Fluorescence was measured via a plate reader.

**Figure S5.** Alignment of RsiV from *B. subtilis*, RsiV from *C. difficile*, and LbpA. The σ-factor binding domain is noted. Underlined residues indicate the transmembrane domain.

**Figure S6.** RsiV and LbpA were overexpressed and purified from *E. coli*. Peptidoglycan from *M. lysodeikticus* was combined with 10 μg/ml hen egg white lysozyme and various concentrations of purified RsiV or LbpA. The A450 was monitored every minute for 30 minutes to determine degradation of lysozyme.

**Figure S7.** Contribution of individual genes in the *csfV* operon on lysozyme resistance. A) MICs of in-frame deletions of individual genes (WT, R20291; Δ*lbpA*, UM303). Overnight cultures were sub-cultured and grown to an OD_600_ of 0.3, incubated with 20 μg/ml lysozyme for 5 hrs, then MICs were determined. B) Single genes were exogenously expressed from a xylose-inducible expression vector in a Δ*csfV* operon mutant strain (Δ*csfV* operon EV, GMK174; Δ*csfV* operon *Pxyl-rsiV*, GMK176; Δ*csfV* operon P_*xyl*_-*lbpA*, GMK178). Cultures were grown with 1% xylose to an OD_600_ of 1.0 then MICs were determined.

**Figure S8.** CRISPRi knockdown was used to determine the contribution of *slpA* and the *dltABCD* operon to lysozyme resistance in a wildtype background, 2 different sgRNAs were tested for *dltABCD* and *csfV* (EV, LS134; *slpA*, GMK344; *dltABCD*, GMK345, GMK346; *csfV*, GMK347, GMK348). Overnight cultures grown with 1% xylose were sub-cultured in TY supplemented with 1% xylose and grown to an OD_600_ of 1.0 then MICs were determined.

